# Divergence of discrete- versus continuous-time calculations of the temperature dependence of maximum population growth rate in a disease vector

**DOI:** 10.1101/2024.08.09.607340

**Authors:** Paul J. Huxley, Leah R. Johnson, Lauren J. Cator, Samraat Pawar

## Abstract

1. The temperature dependence of maximal population growth rate (*r*_m_) is key to predicting how organisms respond and adapt to natural and anthropogenic changes in climate. For organisms with complex life histories, discrete-time matrix projection models (MPMs) can be used to calculate temperature-dependent *r*_m_ because they directly capture variation in empirically-observed life-history trait values as well as the time delays inherent in those traits.
2. However, MPM calculations can be laborious and do not capture the continuous nature of time. Temperature-dependent *r*_m_ calculated from more complex approaches based on delay-differential equation and integral projection models are more accurate but are notoriously difficult to parameterise. Ordinary differential equation-based models (ODEMs) offer a relatively tractable alternative of intermediate complexity but it is largely unknown whether ODEM-based calculations and MPMs broadly agree when the effects of time delays and altered juvenile survival trajectories on temperature-dependent *r*_m_ are introduced by environmental variation.
3. Here we investigate differences in the predicted temperature dependence of *r*_m_ obtained from an ODE-based model with those calculated from MPMs using high-resolution temperature- and resource dependent life-history trait data for the globally-distributed disease vector, *Aedes aegypti*.
4. We show that the level of agreement between discrete- and continuous-time representations of temperature-dependent *r*_m_ can vary with resource availability, and is extremely sensitive to how juvenile survival is characterised. This finding suggests that analytic *r*_m_ models can consistently provide comparable *r*_m_ predictions to standard methods except for under severe resource constraints.
5. Our study also suggests that all formulations of the intrinsic growth rate of a population may not be equally accurate for all types of organisms in all situations. Furthermore, this study’s findings raise questions relating to whether existing mathematical models can be used to predict and understand population-level effects of environmental change.

## Introduction

The distribution and abundance of ectotherm populations is strongly shaped by two key components of their ecological niche: environmental temperature and resource availability. Climate change is predicted to have severe impacts on ectotherms through changes in both (Høye et al., 2021; Lawlor et al., 2024). In particular, how these factors affect arthropod populations, the most abundant group of animals on earth (Bar-On et al., 2018), is of significant concern for ecosystem functioning, agriculture, and human health (Costanza et al., 1997; Outhwaite et al., 2022; Pecl et al., 2017; Ryan et al., 2021; Sánchez-Bayo and Wyckhuys, 2019; Skendžić et al., 2021). For example, recent surges in sustained local dengue transmission across Europe have been partially attributed to how ongoing seasonal temperature changes have increased the region’s thermal suitability for *Aedes* mosquitoes (Branda et al., 2023; Nakase et al., 2023; Oliveira et al., 2021).

To understand how variation in environmental temperature and resource availability can influence arthropods, we need their thermal performance curves (TPCs) for maximal population growth rate (*r*_m_)—a population’s inherent capacity to grow in the absence of density-dependent factors—a fundamental measure of population fitness (Amarasekare and Savage, 2012; Birch, 1948; Cole, 1954; Huey and Berrigan, 2001; Pawar et al., 2024). Previous efforts have used four different approaches for calculating temperature-dependent *r*_m_: matrix projection models (MPMs; Caswell, 1989), ordinary differential equation based models (ODEMs; e.g., Winsor, 1932), delay-differential equation based models (DDEMs; e.g., Amarasekare and Coutinho, 2013; Brass et al., 2021), and Integral Projection models (IPMs; Ellner and Rees, 2006).

Among these methods, DDEMs and IPMs may provide more accurate calculations of *r*_m_ than MPMs and ODEMs but they are the most challenging to use—they are notoriously difficult to parameterise and solve, with IPMs, in addition, being computationally costly. Here we focus on discrete-time MPMs and continuous-time ODEM-based approaches for calculating the TPCs of *r*_m_ because arthropod data are often not at the required resolution for use in these more complex models. For example, IPMs require at least one continuous trait measurement such as age or body size. This parameterisation challenge is amplified when the goal is to calculate fitness across environmental gradients. In contrast, MPMs and ODEM-based approaches can work well with relatively limited data; for example, they can still provide powerful insights into population dynamics even if only stage-specific vital rates (e.g., development, survival and fecundity) are known (Huxley et al., 2021, 2022; Pawar et al., 2024).

For these reasons, MPMs and ODEM-based approaches are commonly used to calculate *r*_m_ in organisms with complex life histories, especially when the goal is to incorporate how different life history stages respond to environmental factors such as temperature. For example, MPMs have been used recently to show that variation in resource availability can have profound effects on the temperature dependence of *r*_m_ in the mosquito vector, *Aedes aegypti* (Huxley et al., 2021, 2022).

MPMs can also capture time-delays in life history cycles. For example, matrix columns can be added to incorporate delays introduced by intermittent resource limitation. Despite these strengths, building MPMs to calculate *r*_m_ across environmental gradients can be laborious and, because they operate in discrete time steps, they typically do not adequately capture the continuous variation in trait values over time. ODE-based models provide a relatively tractable method to calculate *r*_m_ when life history data are incomplete and overcome some of the challenges involved with DDEM and IPM parametrisation. And relative to MPMs, they are less laborious to construct relative and do not suffer from the time-discretization issue. However, the consistency of *r*_m_ calculations based on ODEMs across environmental gradients compared to MPMs is largely unknown.

Here, we study the level of agreement between three approaches to calculate the temperature dependence of *r*_m_ under variation in resource availability in the disease vector, *Aedes aegypti*: an analytic approximation to *r*_m_ derived from an ODE-based model, an explicit MPM simulation, and an analytic approximation to the latter. Our main goal is to evaluate the extent to which *r*_m_ calculated from the continuous time stage-structured population model (the analytic *r*_m_ model; herein the AE-L model) matches that calculated from the MPM across environmental gradients.

We expected any differences in results between these approaches to stem from how differently they weigh the contributions of life-history traits. MPMs and its E-L-based *r*_m_ approximation consider discrete stages and time steps, while continuous-time models and their E-L-based *r*_m_ approximation integrate over life stages and time for juvenile mortality and development. For example, Cator et al. (2020) simplified the AE-L model by assuming a fixed mortality rate (an exponentially-declining survivorship curve), which can be integrated analytically to obtain total cumulative juvenile-stage mortality. In contrast, the MPM and its E-L-based *r*_m_ approximation allow for more varied juvenile survival patterns, which in turn affects predicted *r*_m_. We hypothesize that the AE-L model’s survivorship assumption is valid unless juvenile mortality is high, because, as shown by Cator et al. (2020), *r*_m_ is less sensitive to mortality than to development time and fecundity based on a standard stage-structured ODEM.

We show that the AE-L *r*_m_ model consistently yields similar temperature- and resource-dependent *r*_m_ calculations to the discrete-time E-L equation. However, MPM-derived *r*_m_ calculations can diverge from these approaches due to their sensitivity to how juvenile survival trajectories can be shaped by variation in resource availability.

## Materials and Methods

We tested our hypothesis by modelling juvenile mortality in the E-L equation and MPM as either a fixed reduction per time step (“exponential decay”) or as survival until a cut at the transition point (“transitional decay”). The rationale for this approach is based on how the AE-L model effectively assumes that using an exponentially-decaying survival function to integrate over time steps is reasonable because, in the end, it should still approximate the total proportional mortality at the juvenile-to-adult transition point. In contrast, the E-L equation and MPM can have arbitrary patterns allowing for testing of how different characterisations of juvenile survival may influence *r*_m_. However, in spite of such flexibility, juvenile mortality is rarely directly measured for many arthropods, which forces practitioners to make strong assumptions (e.g., exponential or transitional decay) about it’s dependence on time and environmental variation.

### Calculating r*_m_* using the MPM

We use the standard stage-structured MPM (Caswell, 1989) for population change over discrete time steps:

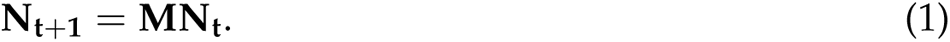

Here, **N_t_** is the vector of abundances in the life stage classes (larval, pupal, and adult) at time *t* and **M** is the transition matrix. The first row of **M** is populated by stage specific daily fecundity (the number of female offspring produced per female at stage/age *i*). The sub-diagonal of **M** is populated with the probabilities of survival from stage *i* to stage *i*+1. Multiplying the transition matrix, **M**, and stage-structured population size vector, **N_t_**, sequentially across time intervals yields the stage-structured population dynamics. Once the stable age/stage distribution (SAD) is reached, the dominant eigenvalue of the system is the finite population rate of increase (*λ*) (Caswell, 1989). A population reaches SAD when the relative proportion of individuals in each life stage has stabilised (that is, the proportion of the population in each age-stage class remains constant over time), even when the total population size is changing (Caswell, 1989). Then, the intrinsic rate of population growth is *r*_m_ = log(*λ*) (i.e., *per capita* change in the population per unit time).

### Approximating r*_m_* from the MPM

Assuming SAD, the discrete-time Euler-Lotka approximation (Lotka, 1907) of the above MPM (Eqn 1) is,

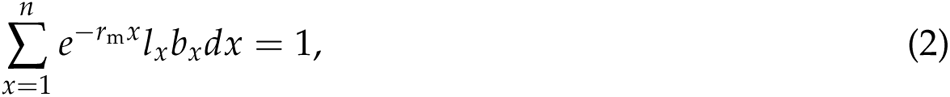

Here, *l_x_* is age/stage-specific survivorship (probability that an individual survives to age/stage *x*), and *b_x_*is stage-specific fecundity (zero for all juvenile stages). This E-L equation describes the expected lifetime reproductive success of a new-born individual in a stage-structured population growing at the rate *r*_m_ once it has reached SAD. To approximate *r*_m_ in Eqn 2, we obtain its root through numerical solution, which requires life tables comprised of *l_x_* and *b_x_* arranged into rows representing age that, in our case, increases daily.

### Calculating r*_m_* from the continuous-time stage-structured population growth model

Analogous to Eqn. 2, under the same assumption of SAD, the E-L equation for the continuous time (ODE-based) model for a age-structured population change is:

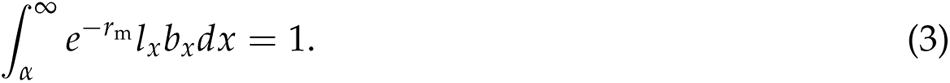

As in Eqn 2, here *r*_m_ is the maximal growth rate once the population has converged on its SAD. However, the life-history traits here are defined slightly differently: *l_x_*and *b_x_* are age-(not stage-) specific survivorship and fecundity, and we now we have a single parameter *α* representing the age of first reproduction corresponding to the development time from egg (or neonate) to reproductive adult. Thus, in this formulation we ignore the discreteness of life stages, a simplification that allows us to derive a closed-form analytic approximation for *r*_m_ (Cator et al., 2020):

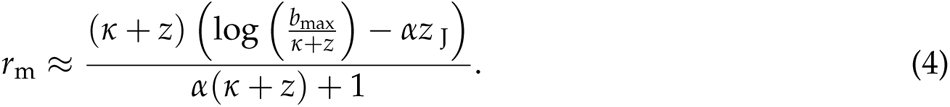

Here, *α* is egg-to-adult development time (days), *b*_max_ is peak reproductive rate (individuals (eggs) *×* individuals (females) *×* day^-1^), *κ* is the fecundity loss schedule (individual^-1^ day^-1^), and *z*_J_ and *z* are juvenile and adult mortality rates (individual^-1^ day^-1^), respectively. Although Eqn 4 is an approximation, it can be shown to be sufficiently accurate as long as *r*_m_ is less than 1 (in units of day^-1^; Cator et al., 2020), which is typically true for insect growth rates (Frazier et al., 2006; Pawar et al., 2024). Equation 4 explicitly incorporates the traits underlying *r*_m_ (through *l_x_* and *b_x_*in Eqn 3), so it can be used to analytically understand how variation in these traits propagates through the system to affect *r*_m_ (Cator et al., 2020). Note that because Eqn 4 is derived from Eqn 3, it is only valid when the population is at SAD, similarly to the MPM and its analytic approximation (Eqn 2).

### Data

To determine the level of agreement between the AE-L model (Eqn 4), the E-L equation and the MPM across environmental gradients, we used two datasets that describe how resource availability can affect *r*_m_ in the disease vector, *Aedes aegypti*. Both datasets are derived from laboratory experiments on this species that were conducted by Huxley et al. (2021; 2022). Huxley et al. (2022) measured the effect of larval competition on the temperature dependence of juvenile development time and mortality rate; *α* and *z*_J_ in Eqn 4, respectively) and adult fitness traits (fecundity rate and mortality rate; *b*_max_ and *z* in Eqn 4, respectively) at five constant temperatures (22, 26, 32, 34 and 36*^◦^*C) and four resource concentration levels (0.183, 0.367, 0.550, and 0.733 mg ml^-1^). Huxley et al. (2021) measured the effect of variable resource supply on the temperature dependence of the same fitness traits at three constant temperatures (22, 26, 32*^◦^*C) and two resource supply levels (0.1 and 1 mg larva^-1^ day^-1^).

### Parametrisation of the r*_m_* models

We parameterised the three *r*_m_ models (Eqns 2, 3, 4) with mean trait responses (development time and adult longevity were rounded to zero decimal places) that we calculated using the raw replicate-level data (*n*=3 replicates at each treatment level; Tables SI1, SI5) from Huxley et al. (2021; 2022). For both datasets, we obtained *z*_J_ by dividing the proportion of juveniles that did not survive to adulthood by *α*. For *z*, we inverted adult longevity (i.e., *z* = 1/longevity), and, as *κ* has been shown to only make a very small contribution to *r*_m_ (Cator et al., 2020), we assumed that *b*_max_ declined with age at a constant rate of 0.01 individual^-1^ day^-1^.

For both datasets, we calculated *r*_m_ for every treatment’s replicates by parametrising the *r*_m_ models with the trait values shown in tables SI1 and SI5. To calculate *r*_m_ with the AE-L model, these values were used directly for each experimental temperature. Due to structural differences between discrete- and continuous-time models, it was necessary to transform these values to estimate *r*_m_ using the E-L equation and the MPM.

Calculating *r*_m_ using the E-L equation requires life tables comprising of rows populated with *l_x_* (the probability that an individual survives to stage *x*) and *b_x_* (stage-specific fecundity; set to zero for all juvenile stages). The total number of rows in each life table was equal to the sum of development time plus adult longevity. At maturity, *b_x_* was equal to *b*_max_, which decreased at a rate of *κ* per time step (*b*_max_ *−* (0.01 *×* day) until death, i.e., the last row of the life table). When juvenile survival probability (*l_x_*) was assumed to decrease at a fixed rate per day, this quantity was obtained by subtracting *z*_J_ *×* developmental day from 1. When juvenile survival only decreased at the transition point to adulthood, life table rows were populated with 1 until this point. At transition, juvenile survival was obtained by subtracting *z*_J_ *×* total development time (i.e., *α*) from 1. Adult survival decreased at a fixed rate per day in all life tables. This quantity was obtained by subtracting 1/adult longevity *×* day from the juvenile survival probability at transition until the final adult age class (i.e., row).

To estimate *r*_m_ using the MPM, each column in the transition matrix (**M** in Eqn 1) represented one day. The total number of columns in **M** was equal to the sum of development time plus adult longevity. Reproduction does not occur in the juvenile stages, so the first row of **M** was populated with zeros until maturity. At maturity, the first row of **M** was populated with *b*_max_ which decreased at a rate of *κ* per time step (i.e., *b*_max_*−*(0.01 *×* day)) until death. The sub-diagonal of **M** was populated with the probabilities of survival from stage *t* to stage *t*+1. As with the life tables for the E-L equation, juvenile survival **M** was obtained by subtracting *z*_J_ *×* day from 1. When juvenile survival decreased at the transition point to adulthood, the sub-diagonal was populated with 1 until this point. At transition, juvenile survival was obtained by subtracting *z*_J_ *×* total development time (i.e., *α*) from 1. Adult survival was obtained by subtracting 1/adult longevity *×* day from the juvenile survival probability at transition until the final adult age class. The projection matrices were built and analysed in R (R Core Team, 2023) using the popbio package (Stubben and Milligan, 2007). To estimate *r*_m_ for the E-L equation, we used the uniroot function in base R (R Core Team, 2023).

### Fitting Thermal Performance Curves (TPCs) to the r*_m_* calculations

The temperature at which *r*_m_ peaks (*r*_m_ *T*_opt_) and the value of *r*_m_ at its peak (*r*_opt_) are important parameters for understanding how arthropod populations will respond to long-term sustained climatic warming (Pawar et al., 2024). To predict *r*_m_ *T*_opt_ and *r*_opt_, we generated continuous *r*_m_ TPCs using non-linear least squares (NLS) in the rTPC pipeline (Padfield et al., 2021). We fitted the Sharpe-Schoolfield TPC model (Kontopoulos et al., 2018; Schoolfield et al., 1981; Eqn SI1) to the replicate-level *r*_m_ calculations for each experimental temperature in the larval competition dataset (Huxley et al., 2022). We used this model because it has been theoretically and empirically validated. However, as is the case with most TPC models, the Sharpe-Schoolfield model (Schoolfield et al., 1981) can only be used to predict non-negative responses, so for treatments comprising of both non-negative and negative *r*_m_ calculations, we fitted a generalised additive model (GAM) using the mgcv package (Wood and Wood, 2015; Wood, 2011).

The resource limitation dataset (Huxley et al., 2021) does not cover a sufficient range of temperatures to fit continuous *r*_m_ TPCs, so we parametrised Eqn 4 with the data for each temperature *×* resource supply treatment in that study.

## Results

### Comparison of the r*_m_* calculations using the larval competition dataset

At all resource levels in the larval competition dataset (Huxley et al., 2022), the AE-L model and the E-L equation predicted *r*_m_ to respond unimodally to temperature — it increased as temperatures increased from 22 to 32–34*^◦^*C before declining rapidly to negative growth (figure 1). For both characterisations of juvenile survival, the *r*_m_ calculations for the AE-L model and the E-L equation increased from *∼*0.1 at 22*^◦^*C to *∼*0.2 at 34*^◦^*C at resource levels above 0.183 mg ml*^−^*^1^.

**Figure 1:**
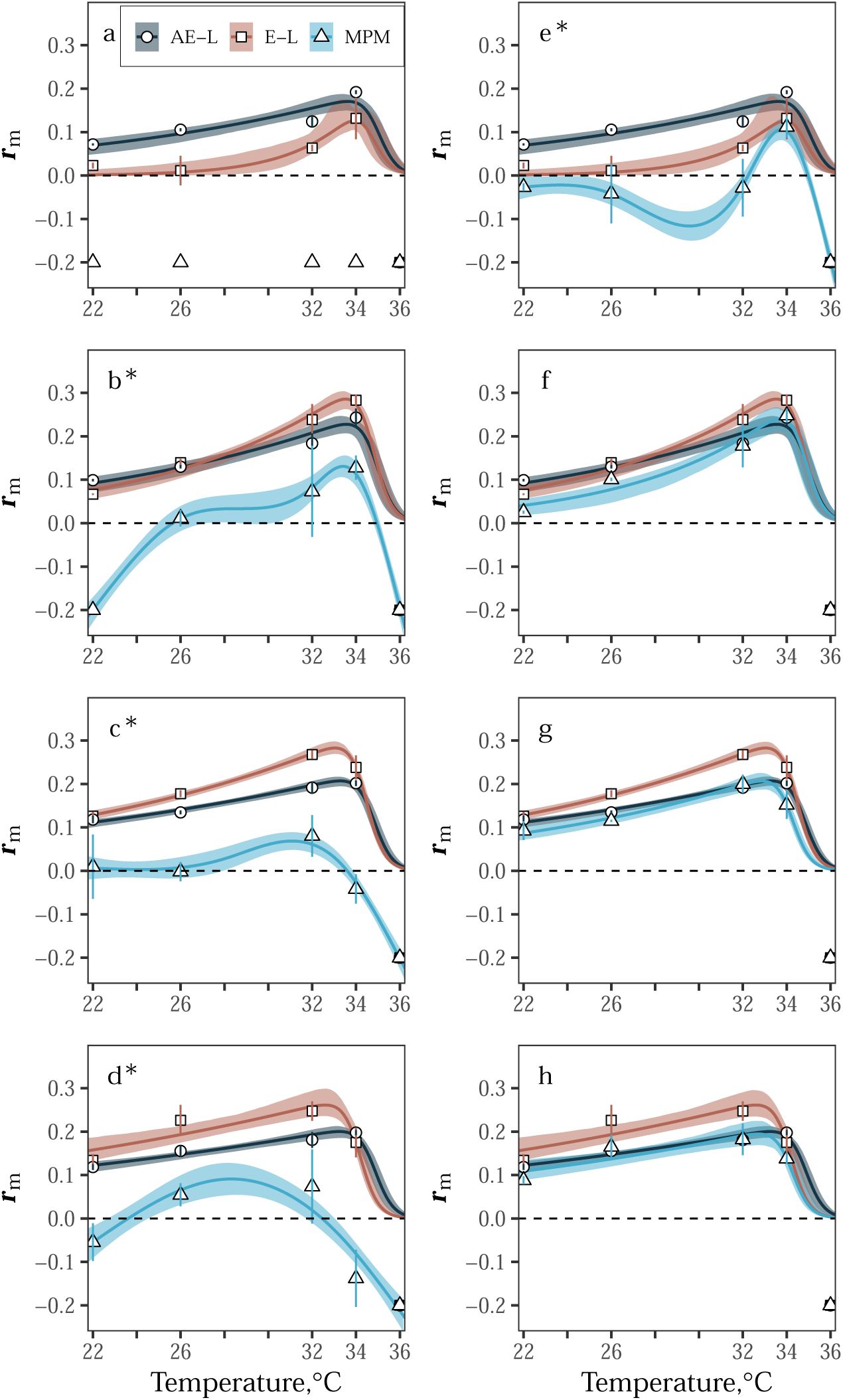
TPCs fitted to *r*_m_ calculations from the analytic *r*_m_ model (AE-L, grey), the Euler-Lotka equation (E-L, red) and the MPM (blue) for the resource competition dataset (Huxley et al., 2022). (a – d) *r*_m_ TPCs for the three *r*_m_ models across resource concentration levels (0.187, 0.367, 0.550, and 0.733 mg ml^-1^, respectively). For the E-L equation and the MPM, juvenile survival decreased at a fixed rate per time step. (e – h) *r*_m_ TPCs at the same resource levels as (a – d) but for the E-L equation and the MPM, juvenile survival only decreased at the juvenile-to-adult transition point. Symbols denote mean *r*_m_ *±* standard deviation (error bars) calculated from the *r*_m_ calculations for each treatment’s replicates (*n*=3; replicate-level *r*_m_ calculations are provided in the manuscript’s accompanying GitHub repository). *r*_m_ TPCs were fitted to Eqn SI1 using rTPC (Padfield et al., 2021). Asterisks (*) indicate TPCs (blue) fitted to MPM-derived *r*_m_ calculations (triangles) using GAMs. Bootstrapping (residual resampling) was used to calculate 95% prediction bounds for each *r*_m_ TPC. Negative *r*_m_ calculations were cut off at –0.2 for plotting.

In contrast, the *r*_m_ calculations from the AE-L model differed from the E-L equation derived *r*_m_ calculations at 0.183 mg ml*^−^*^1^. The AE-L model predicted *r*_m_ to increase from 0.07 at 22*^◦^*C to 0.19 at 34*^◦^*C, whereas the E-L equation predicted *r*_m_ to increase from 0.02 to 0.13 across these temperatures. Under transitional decay at higher resource levels (*>*0.183 mg ml*^−^*^1^), the MPM-derived *r*_m_ calculations were generally consistent with the *r*_m_ calculations from the A-EL model and the E-L equation (figure 1e–h), but, at 0.183 mg ml*^−^*^1^, MPM-derived *r*_m_ was only positive at 34*^◦^*C (figure 1e).

At all resource levels, the MPM-derived *r*_m_ calculations differed from the other approaches when juvenile survival decayed exponentially. Under this assumption at 0.183 mg ml*^−^*^1^, MPM-derived *r*_m_ was negative at all temperatures, and, at higher resource levels MPM-derived *r*_m_ was negative or close to zero at temperatures below 26*^◦^*C. MPM-derived *r*_m_ increased above this temperature, but it became negative again at 34*^◦^*C or peaked at lower value than it did for the AE-L model and the E-L equation (figure 1c & d).

Except for the MPM TPC at 0.733 mg ml*^−^*^1^ under exponential decay, the TPCs for all three *r*_m_ models predicted *r*_m_ to peak (*T*_opt_) between 31–34*^◦^*C, irrespective of juvenile survival assumption or resource level (figure 2, table SI4). Optimal thermal fitness (*r*_opt_) was, in most cases, consistent across resource levels and survival assumptions for all *r*_m_ models. It was not possible to predict *r*_opt_ under exponential decay for the MPM because *r*_m_ was negative at all temperatures. However, at 0.183 mg ml*^−^*^1^ under transitional decay, MPM *r*_opt_ was predicted to be lower than *r*_opt_ for both the AE-L model (0.11 compared to 0.17, respectively), and the E-L equation (0.13). At all resource levels under exponential decay, MPM *r*_opt_ was predicted to be lower than MPM *r*_opt_ under transitional decay. The *r*_opt_ predictions for the MPM under exponential decay were also lower than the *r*_opt_ predictions for the AE-L model and the E-L equation, as noted above (figure 2, table SI4).

**Figure 2:**
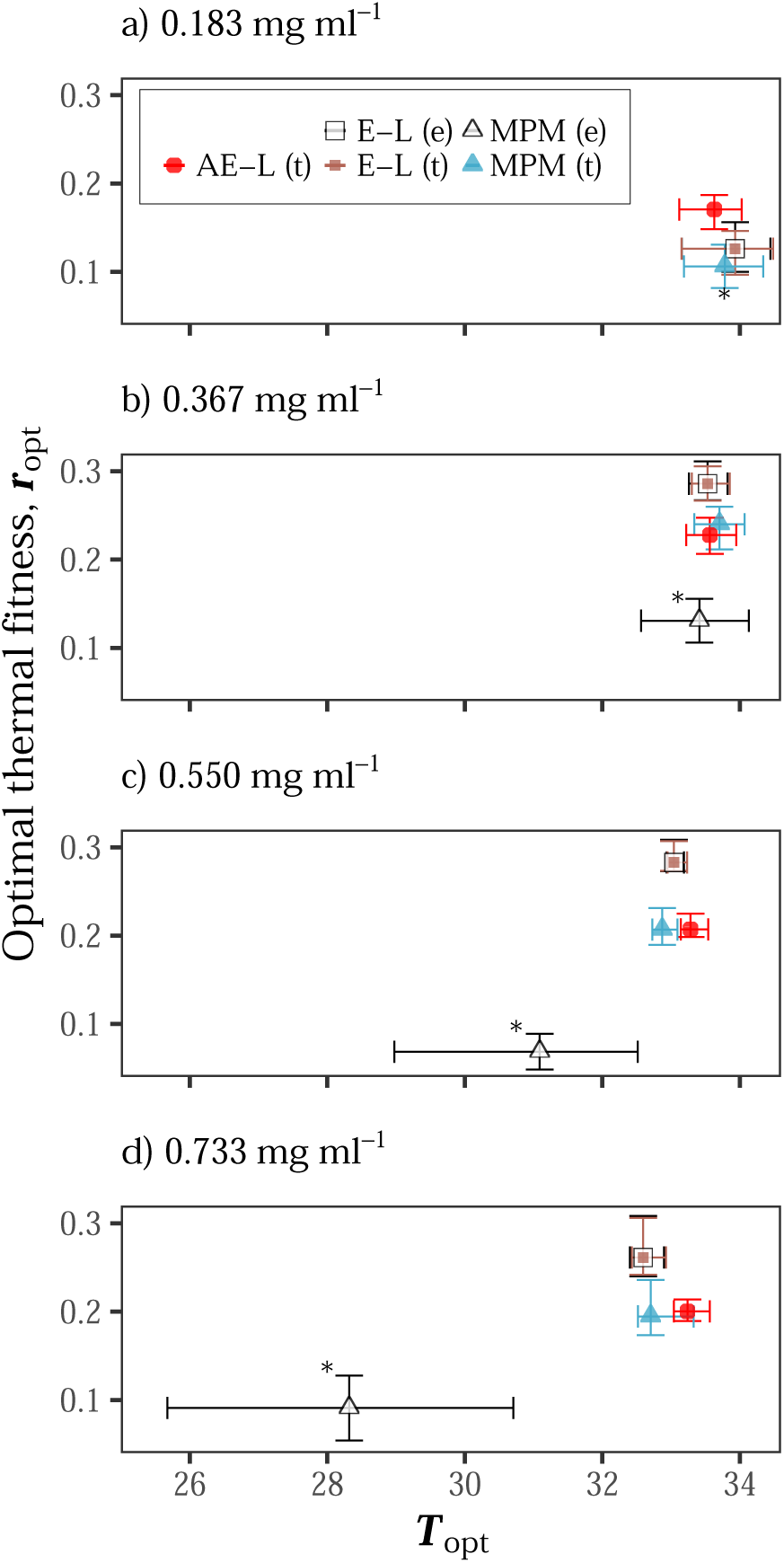
Comparison of predicted optimal thermal fitness (*r*_opt_) at *T*_opt_ for the analytic *r*_m_ model (AE-L), the Euler-Lotka equation (E-L), and the MPM for the resource competition dataset (Huxley et al., 2022). **a-d** Predicted *r*_opt_ at *T*_opt_ for the three *r*_m_ models across increasing larval resource concentration levels. Predictions (medians) with bi-directional error bars for when juvenile survival decreased at a fixed rate per time step are denoted “(e)” in the legend. Predictions for when juvenile survival only decreased at the juvenile-to-adult transition point are denoted “(t)”. Points with asterisks (*) are *r*_opt_s from TPCs fitted to MPM-derived *r*_m_ calculations using GAMs. Bootstrapping (residual resampling) was used to calculate 95% uncertainty bounds for each point.

### Comparison of the r*_m_* calculations using the larval resource supply dataset

Under both survival assumptions at high-resource supply (1 mg larva*^−^*^1^ day*^−^*^1^) in the resource limitation dataset (Huxley et al., 2021), all *r*_m_ models predicted *r*_m_ to be positive and increase monotonically with temperature to its peak at 32*^◦^*C. Under both juvenile survival assumptions at high-resource supply, differences between the AE-L and the E-L were small (figure 3ab, table SI6), but MPM *r*_m_ under exponential decay was lower than *r*_m_ from the AE-L model and the E-L (figure 3a, table SI6).

**Figure 3:**
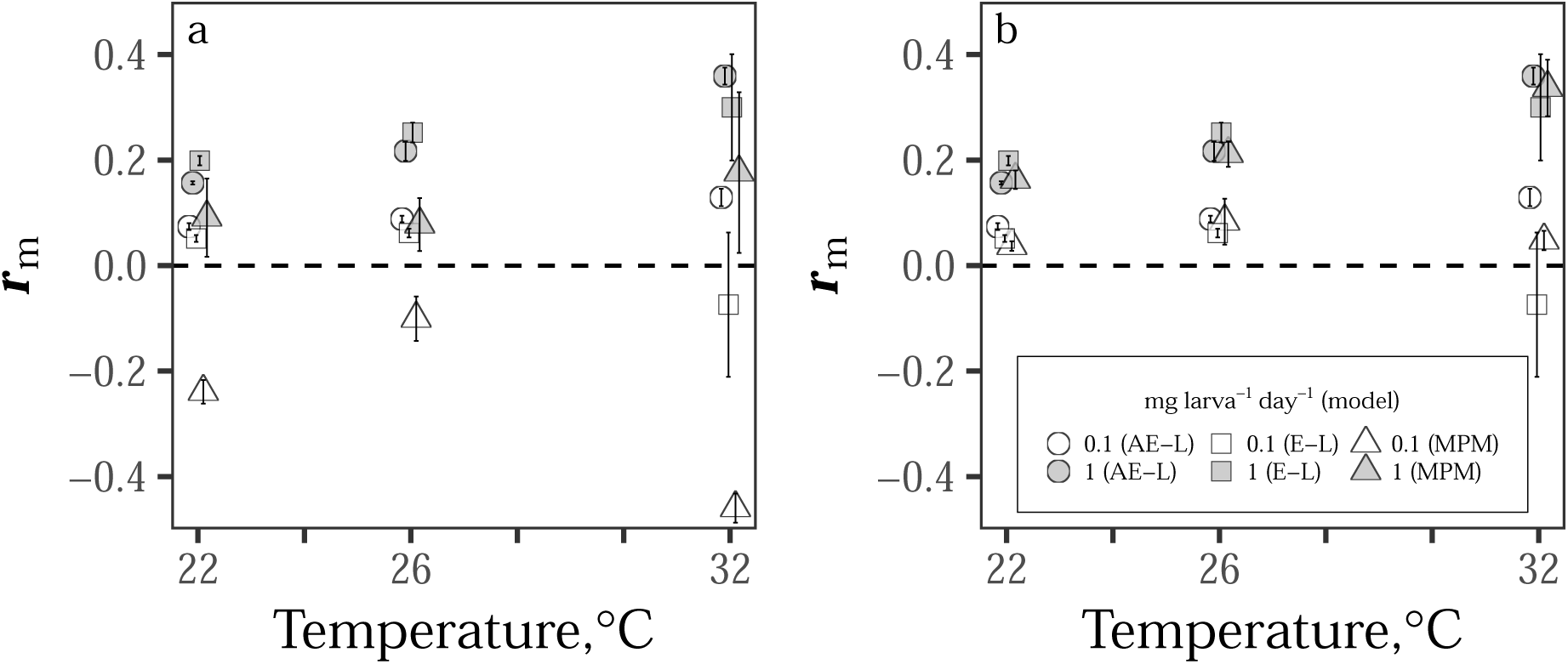
Comparison of *r*_m_ calculations from discrete- and continuous-time *r*_m_ models for the resource limitation dataset (Huxley et al., 2021). Estimated *r*_m_ from the analytic *r*_m_ model (AE-L), the Euler-Lotka equation (E-L), and the MPM at two larval resource supply levels (0.1 and 1 mg larva^-1^ day^-1^, white and grey symbols, respectively). **(a)** Calculations under the assumption that juvenile survival decreases at a fixed rate per time step. **(b)** Calculations when juvenile survival is assumed to only decrease at the juvenile-to-adult transition. Symbols denote mean *r*_m_ *±* standard deviation (error bars) calculated from the *r*_m_ calculations for each treatment’s replicate populations (*n*=3; replicate-level *r*_m_ calculations are provided in the manuscript’s accompanying GitHub repository.)

At low-resource supply (0.1 mg larva*^−^*^1^ day*^−^*^1^) under the exponential decay assumption in the larval resource supply dataset, the MPM *r*_m_ calculations were always negative (figure 3a, table SI6). In contrast, AE-L model *r*_m_ was always positive, and the E-L equation *r*_m_ calculations were positive at 22 and 26*^◦^*C before becoming negative at 32*^◦^*C. The MPM predicted *r*_m_ to increase from –0.24 at 22*^◦^*C to –0.10 at 26*^◦^*C; it then decreased to –0.46 at 32*^◦^*C. The AE-L model predicted *r*_m_ to increase with temperature from 0.07 at 22*^◦^*C to 0.09 at 26*^◦^*C, and 0.13 at 32*^◦^*C, and the E-L equation predicted *r*_m_ to increase from 0.05 at 22*^◦^*C to 0.06 at 26*^◦^*C and then decrease to –0.07 at 32*^◦^*C.

## Discussion

Our goal was to examine the extent to which an analytic *r*_m_ model based on the continuous E-L equation (the AE-L model; Eqn 4) can provide similar calculations of *r*_m_ when parameterized with data across resource availability gradients, in comparison with that calculated from the (discrete) E-L equation (Eqn 2) and the MPM (Eqn 1).

Using two datasets that describe how resource availability and temperature jointly affect traits, and thus the predicted temperature dependence of *r*_m_ (Huxley et al., 2021, 2022), we show that the AE-L model consistently yields similar calculations to the discrete time E-L equation when populations are not constrained by larval resource limitation or competition effects. This is as we expect, since the AE-L is based on the continuous time version of the E-L equation as we explain further below. We also show that modellers should proceed with caution when resource limitation or competition effects are expected to strongly mediate the temperature dependence of *r*_m_ as model predictions differ in these cases.

For both datasets (Huxley et al., 2021, 2022), when resource availability was high and juvenile survival was assumed to only decrease at the juvenile-to-adult transition point, the AE-L model was also consistently in agreement with the MPM *r*_m_ calculations across temperatures. Under transitional decay, the margin of error between all sets of *r*_m_ calculations for the high resource treatments was generally small (figures 1f-h, 3b; tables SI2, SI3, SI6).

Under transitional decay at resource levels higher than 0.183 mg ml*^−^*^1^ in the larval competition dataset (Huxley et al., 2022), all approaches predicted *r*_m_ to respond unimodally to temperature — increasing with temperature from 22 to *∼*33*^◦^*C, and then declining rapidly to zero after this peak (figures 1, 2; tables SI2–SI4). Similarly, for the high-resource supply (1 mg larva*^−^*^1^ day*^−^*^1^) in the resource limitation dataset (Huxley et al., 2021), the AE-L model consistently agreed with the *r*_m_ calculations from the E-L equation and the MPM (i.e., again, the margin of error between the three sets of *r*_m_ calculations for these treatments was generally not substantial; figure 3c, table SI6). For the high-resource supply treatments in the resource limitation dataset (Huxley et al., 2021) and for both survival assumptions, all approaches (with the exception of the small decrease in *r*_m_ as temperatures increased from 22 to 26*^◦^*C for the MPM) predicted *r*_m_ to be positive and increase monotonically as temperatures increased from 22 to 32*^◦^*C.

In contrast, for both datasets (Huxley et al., 2021, 2022) when resource availability was low, the AE-L model calculations and the MPM *r*_m_ calculations markedly diverged (figures 1ab & 3a, tables SI2, SI3 & SI6). The largest discrepancy between the three approaches in the larval competition dataset (Huxley et al., 2022) was at the lowest resource level (0.183 mg ml*^−^*^1^ day*^−^*^1^) under the assumption that juvenile survival decreased by a fixed rate per time step. It was not possible to estimate *T*_opt_ and *r*_opt_ for the MPM under this assumption because *r*_m_ was negative across all of the temperature range, yet both the AE-L model and the E-L equation predicted *T*_opt_ to be close to 34*^◦^*C and *r*_opt_ to be *∼*0.15. There were also important differences between the *r*_m_ models for the low resource level under transitional decay of juvenile survival. The three approaches predicted *T*_opt_ to be close to 34*^◦^*C and *r*_opt_ to be *∼*0.15, but the E-L equation predicted *r*_m_ to be close to zero between 22 and 32*^◦^*C, and the MPM differed in its prediction of the sign of *r*_m_ across much of the temperature range (figure 1e, table SI4).

Under the model assumption of exponentially decaying juvenile survival, an even greater level of disparity was observed between the AE-L prediction and the MPM model for the resource limitation dataset (Huxley et al., 2021) at low-resource supply (0.1 mg larva*^−^*^1^ day*^−^*^1^). In this case, the AE-L model predicted *r*_m_ to be positive (and increasing) across the entire temperature range, whereas as the MPM predicted *r*_m_ to be negative across this range. The E-L *r*_m_ prediction was intermediate between them (figure 3a, table SI6).

This disagreement in the calculations of *r*_m_ amongst the models at low resource levels (Huxley et al., 2021, 2022) stems from differences in their mathematical structure. Stage-structured MPMs project population abundance over discrete time intervals, assume discrete stages, and (as formulated here) explicitly consider all juvenile stages. In contrast, the continuous-time AE-L model aggregates these stages into a single, continuous stage described by a single, cumulative juvenile mortality and development time parameter (*z*_J_ and *α*, respectively). This simplifying assumption of the AE-L model appears to affect the calculations of *r*_m_ when populations are exposed to environmental conditions that cause delays in the system.

Also, any differences between the *r*_m_ models must stem from the aggregation of juvenile mortality and development in the AE-L model because we have essentially compared *r*_m_ calculated from the discrete-time Euler-Lotka equation (Eqn 2) to an approximation of *r*_m_ derived from the continuous Euler-Lotka equation (Eqn 3). Essentially, the aggregation of juvenile mortality and development in the AE-L model makes it relatively insensitive to juvenile survival. While the discrete-time E-L equation is more sensitive to proportional juvenile survival at transition, the fact that its *r*_m_ calculations were the same irrespective of how survival decreased prior to the juvenile-to-adult transition point shows that it is insensitive to the specifics of how juvenile survival is characterised. The MPM, on the other hand, is highly sensitive (perhaps too sensitive) to how juvenile survival is characterised. This key finding shows that standard practices for characterising juvenile survival, whether that be integrating over time steps (in the case of the AE-L) or assuming a constant rate of mortality (i.e., MPMs) are problematic, particularly when resources are limiting, and as a result may be age or size specific. Indeed, different juvenile survival characterisations can also introduce substantial bias when converting age-structured vital rates estimated from life tables to calculate *r*_m_ using MPMs that are only stage-structured (Fujiwara and Diaz-Lopez, 2017; Kendall et al., 2019).

Environmental differences that cause time delays in life history can be directly accounted for in MPMs by increasing the number of columns in the transition matrix (**M**, Eqn 1) assigned to a particular life stage. For example, intensified larval competition at low resource levels increases development time across all juvenile stages (Huxley et al., 2022). This effect can be accounted for in MPMs by increasing the number of columns in **M** assigned to the juvenile life stages. In this way, MPMs implicitly include time delays. This implicit “stretching” of **M** to account for delays makes it possible to study how environment-driven delays affect *r*_m_. Delay effects can also be studied using continuous-time stage structured population models (Gurney et al., 1983; Nisbet and Gurney, 1983) that introduce delays explicitly with delay parameters (thus yielding delay-differential equations). In contrast, by merging the juvenile stages into a single, continuous stage, the AE-L model may not adequately weight the negative impact that delay mechanisms can have on *r*_m_ since these factors are effectively summarized in a single parameter representing the expected time of maturation. This simplifying assumption of the analytic *r*_m_ model implies that delay mechanisms are expected to only have negligible effects on *r*_m_.

The results of the MPM sensitivity analyses in Huxley et al. (2021, 2022) are qualitatively similar with the sensitivity analysis of the AE-L model in Cator et al. (2020) in showing that juvenile traits contribute more to *r*_m_ than adult traits. This also supports the notion that the AE-L may not reliably estimate *r*_m_ in low resource conditions because it does not adequately account for the negative effect of increased juvenile mortality on *r*_m_. Additional examination of these sensitivity analyses provides further insights into the behavior of the AE-L model’s estimate of *r*_m_ in low resource conditions. For example, the sensitivity analysis for the AE-L model indicates that juvenile survival contributes a smaller proportion to *r*_m_ than indicated by the MPM sensitivity analysis and this model’s *r*_m_ calculations shown here. While the MPM sensitivity analyses in Huxley et al. (2021, 2022) indicate that together juvenile development time and survival contribute more to *r*_m_ than adult traits do, their respective contributions *r*_m_ cannot be easily separated. However, when the MPM sensitivity analyses in Huxley et al. (2021, 2022) are used in combination with the MPM *r*_m_ calculations reported here at low resource levels, it is clear that MPM *r*_m_ is more sensitive to juvenile survival than development time. For example, the number of matrix columns assigned to the juvenile stages (i.e., development time) were identical for the low resource supply level MPMs in Huxley et al. (2021), yet under exponentially decaying survival the MPM predicted *r*_m_ to be –0.24 compared to 0.04 under transitional decay.

The first implication of this finding is that not all of the formulations of the intrinsic growth rate of a population may be equally accurate for all types of organisms in all situations. As we find here, for organisms where juvenile survival patterns are neither constant nor exponentially distributed (either generally or due to environmental conditions) the choice of metric matters to conclusions drawn about the population. Thus either an MPM approach may be preferred (when a discretization is appropriate) or when a continuous time model is preferred, a recognition that the calculated *r*_m_ may be overly optimistic should be kept in mind. Although the analytic AE-L model may be unreliable at low resource levels, a key finding of this study — that the AE-L model can reliably predict *r*_m_ when populations are not constrained by resource limitation or larval competition effects — has important implications for the efficiency of study workflows. For example, this method allows for much easier integration of laboratory measures of trait performance to investigate the effect of temperature at high resource supply than individually constructing MPMs for each treatment. Furthermore, with respect to VBD transmission frameworks specifically, this result shows that continuous-time analytic *r*_m_ models may offer a simple method for “plugging in” *r*_m_ responses into continuous-time VBD models. Integrating these models into broader VBD model frameworks could improve their reliability in predicting of how temperature and high resource levels together affect transmission risk through their effects on vector *r*_m_.

The second important implication of this study’s central finding relates to whether existing mathematical models can be used to predict and understand the population-level effects of environmental change. The answer to this question lies with two connected ideas. First, whether under temperature fluctuations alone any of these approaches capture meaningful summaries of population performance either in aggregate or instantaneously. Currently, calculations of *r*_m_ at fixed temperatures are often averaged or used for rate summation-type approaches in order to estimate a realized/time-averaged *r*_m_. However, we know of no laboratory experiments that confirm multi-generational population growth rates under known fluctuation regimes are well captured by the approach (although in other contexts the generalization from constant to varying temperature is known to be fraught). Further, whether these metrics are accurate (or useful) measures of temperature-dependent *r*_m_ in the field and how they are affected by other environmental factors, such as resource availability must be assessed. For example, existing model frameworks (e.g., continuous-time stage-structured population models; Amarasekare and Coutinho, 2013; Beck-Johnson et al., 2013) can be developed to incorporate temperature- and resource-induced developmental delays, but datasets that describe these combined effects in the field are largely absent. The absence of such datasets is probably due to the fact that new measures are needed to determine how effective temperature-dependent *r*_m_ in the field is affected by resource fluctuations. Further, realistic and tractable measures of density-dependent effects on *r*_m_ are needed to be able for predicting the effects of environmental change on insect abundance dynamics, in general.

Semi-field systems could provide opportunities to track the entire insect life cycle under ambient environmental conditions. For example, in *Drosophila*, such systems have recently been used to observe thermal adaptation in response to natural environmental change by tracking the evolution of fitness-associated phenotypes and allele frequencies (Rudman et al., 2022) across generations. In mosquitoes, such systems have generally been used to test the effectiveness of novel bio-control strategies, such as transgenic fungi (Lovett et al., 2019), but they also could allow for the effects of temperature *×* resource interactions in the larval stage on fitness and abundance to be explored under conditions which more closely resemble natural environments. Further insights could be provided by examining the interaction between insects and microbes, for example. Recent studies show that mosquitoes can be reared exclusively on cultures of *Asaia* bacteria (Chouaia et al., 2012; Souza et al., 2019), while other studies have shown that larval exposure to microbial variability can affect adult mosquito life history traits (Dickson et al., 2017). However, microbiota at mosquito breeding sites is spatially heterogeneous (Hery et al., 2021), which could mean that any generalizable patterns in resource availability could be difficult to detect.

In spite of the difficulties posed by this challenge, greater research effort towards this important issue is needed, especially if the goal is to understand how insects, including vector populations and VBD transmission patterns, will respond to climatic warming. Indeed, resource availability itself is likely to be temperature-dependent because microbial growth rates also increase with temperature (Craine et al., 2010; Cross et al., 2015; Smith et al., 2019). In this way, increases in environmental temperatures could increase the concentration of food in the environment, which will increase population growth through decreased juvenile development time and increased adult recruitment rates. This could contribute to the expansion of disease vectors and other invasive insect species into regions that were previously prohibitive by broadening *r*_m_’s thermal niche width (Amarasekare and Simon, 2020; Huxley et al., 2021, 2022; Lehmann et al., 2020).

Recent studies have used *r*_m_ calculations from the Euler-Lotka equation to predict population viability under climatic warming (Deutsch et al., 2008; Tewksbury et al., 2008), but very few studies have assessed whether the Euler-Lotka equation can provide reliable estimates of *r*_m_ when populations are exposed to multiple environmental factors. Our study shows that analytic models based on the Euler-Lotka equation can provide similar calculations of temperature-dependent *r*_m_ to standard methods providing resource environments are non-limiting. This study also underlines the need for accurate measures of how variation in resource availability in the field can affect the thermal response of *r*_m_ and therefore, abundance. Such data are particularly key to improving predictions of how climatic warming will affect seasonal insect populations through its effects on abundance dynamics.

## Supporting Information

### Supplementary Equation

#### Modelling the r*_m_* TPCs

We modelled the *r*_m_ TPCs using the modified Sharpe-Schoolfield equation (Equation SI1; Kontopoulos et al., 2018; Schoolfield et al., 1981):

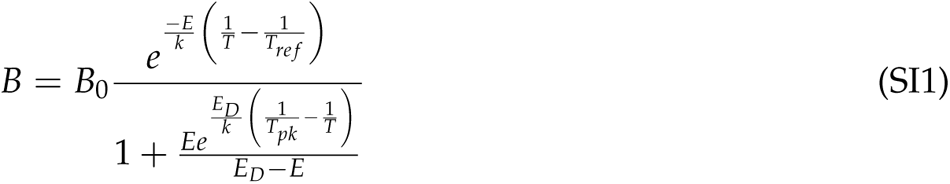

Here, *B* is the value of the metabolic trait at a given temperature (*T* (K)), *B*_0_ is a normalisation constant representing the value of the trait at some reference temperature (*T_re f_*), *E* is the apparent activation energy (initial thermal sensitivity), which determines how fast the curve rises up to *T_pk_*, and *E_D_* is the deactivation energy, which determines how fast the trait declines after the peak. The parameter *k* is the universal Boltzmann constant (8.617 *·* 10*^−^*^5^ eV *K^−^*^1^).

Equation A1 has been used as a model for thermal performance of traits in numerous previous studies on arthropod population biology because it accurately captures the temperature dependence of a wide range of metabolically constrained life history traits.

As is the case with most TPC models, the Sharpe-Schoolfield model (Schoolfield et al., 1981) can only be used to predict non-negative responses, so we fitted generalized additive models (GAM), with the mgcv R package (Wood and Wood, 2015), using restricted maximum likelihood (REML; Wood, 2011) to fit TPCs for treatments comprising of both non-negative and negative *r*_m_ calculations. All data and code to reproduce the results, figures and tables are in an anonymised Github repository.

**Table S1:**
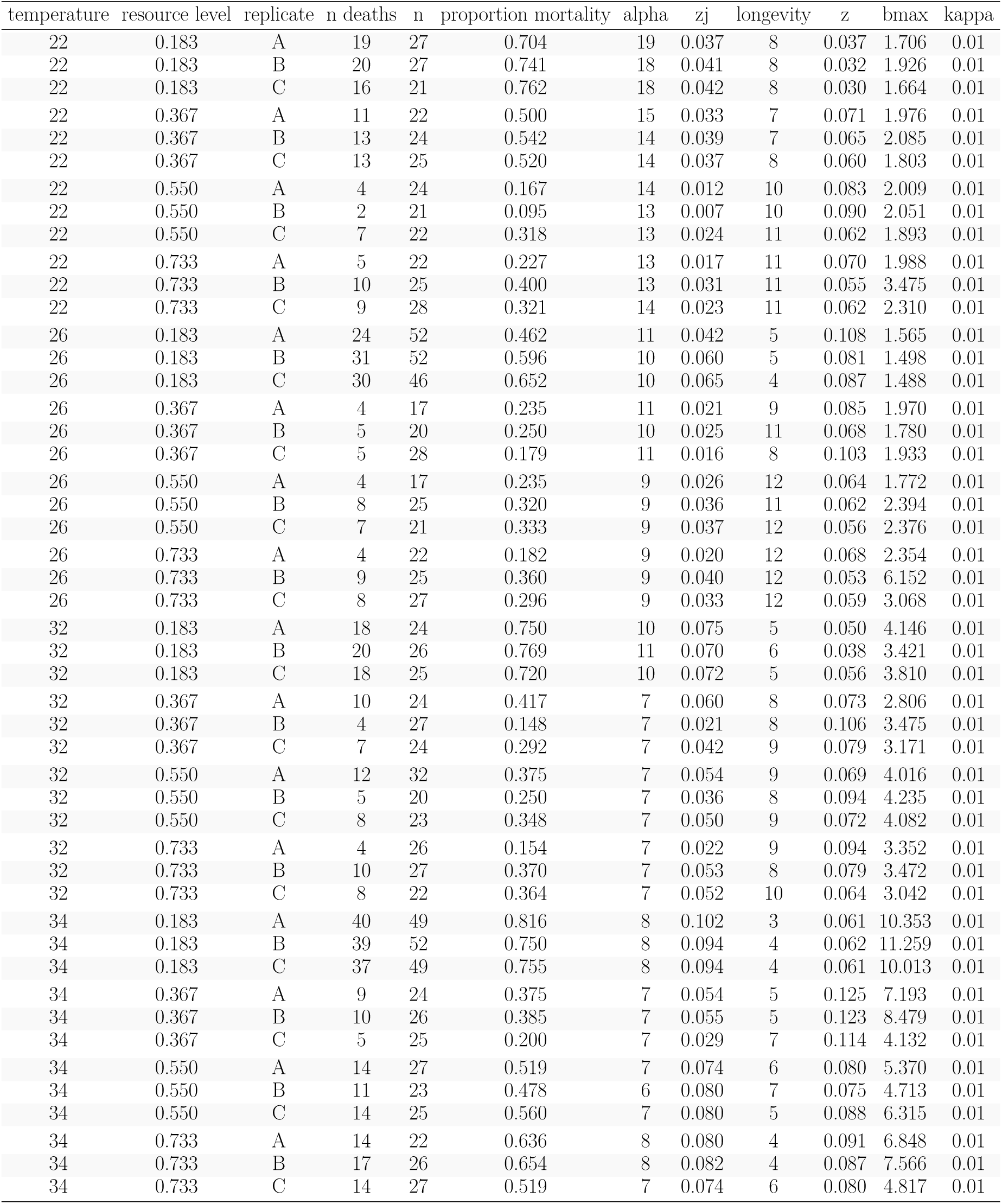
Replicate-level fitness trait values used to calculate *r*_m_ using the analytic *r*_m_ model, the E-L equation, and the MPM for the larval competition dataset (Huxley et al., 2022). *n* is the number of individuals in each replicate at the start of the experiment, *α* is juvenile development time (days), *b*_max_ is peak reproductive rate (individuals (eggs) *×* individuals (females) *×* day^-1^), *κ* is the fecundity loss schedule (individual^-1^ day^-1^), and *z*_J_ and *z* are juvenile and adult mortality rates (individual^-1^ day^-1^), respectively.

**Table S2:**
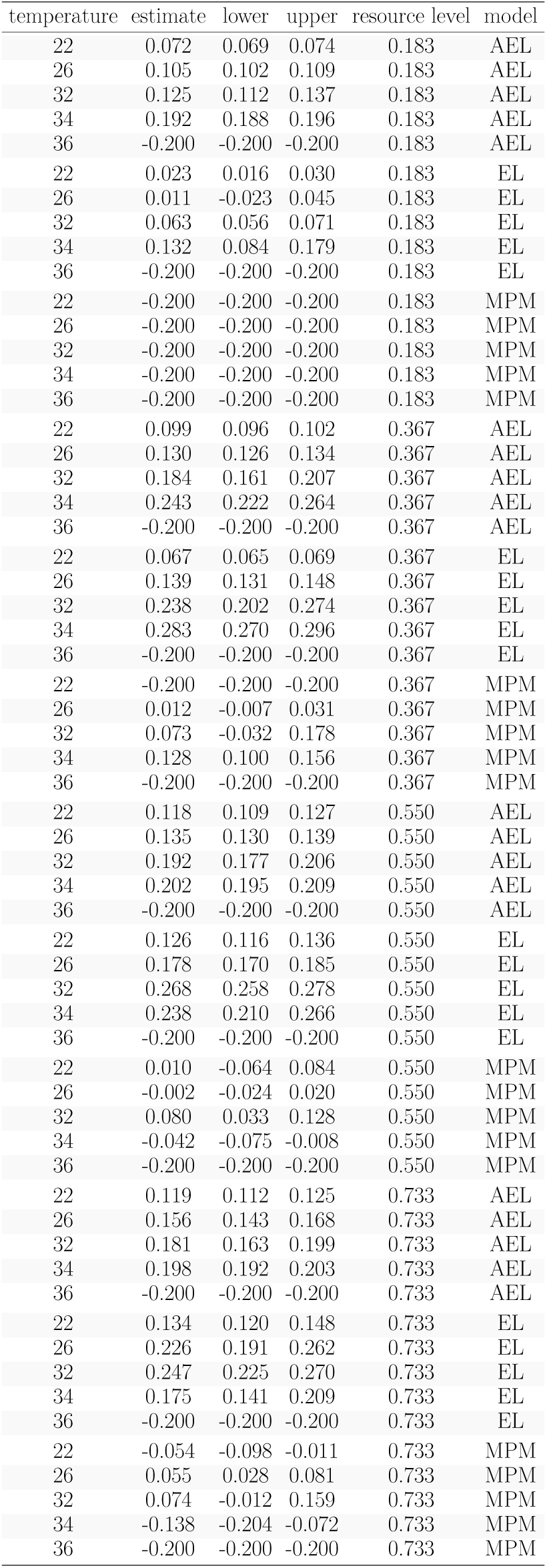
*r*_m_ calculations (mean *±* standard deviation) calculated using the analytic *r*_m_ model (AE-L), the E-L equation, and the MPM for the larval competition dataset (Huxley et al., 2022). For the E-L equation and the MPM, juvenile survival was assumed to decrease at fixed rate per time step. Resource levels are in mg ml*^−^*^1^. Replicate-level *r*_m_ calculations are provided as an accompanying data file.

| temperature | estimate | lower | upper | resource level | model |
| --- | --- | --- | --- | --- | --- |
| 22 | 0.072 | 0.069 | 0.074 | 0.183 | AEL |
| 26 | 0.105 | 0.102 | 0.109 | 0.183 | AEL |
| 32 | 0.125 | 0.112 | 0.137 | 0.183 | AEL |
| 34 | 0.192 | 0.188 | 0.196 | 0.183 | AEL |
| 36 | -0.200 | -0.200 | -0.200 | 0.183 | AEL |
| 22 | 0.023 | 0.016 | 0.030 | 0.183 | EL |
| 26 | 0.011 | -0.023 | 0.045 | 0.183 | EL |
| 32 | 0.063 | 0.056 | 0.071 | 0.183 | EL |
| 34 | 0.132 | 0.084 | 0.179 | 0.183 | EL |
| 36 | -0.200 | -0.200 | -0.200 | 0.183 | EL |
| 22 | -0.200 | -0.200 | -0.200 | 0.183 | MPM |
| 26 | -0.200 | -0.200 | -0.200 | 0.183 | MPM |
| 32 | -0.200 | -0.200 | -0.200 | 0.183 | MPM |
| 34 | -0.200 | -0.200 | -0.200 | 0.183 | MPM |
| 36 | -0.200 | -0.200 | -0.200 | 0.183 | MPM |
| 22 | 0.099 | 0.096 | 0.102 | 0.367 | AEL |
| 26 | 0.130 | 0.126 | 0.134 | 0.367 | AEL |
| 32 | 0.184 | 0.161 | 0.207 | 0.367 | AEL |
| 34 | 0.243 | 0.222 | 0.264 | 0.367 | AEL |
| 36 | -0.200 | -0.200 | -0.200 | 0.367 | AEL |
| 22 | 0.067 | 0.065 | 0.069 | 0.367 | EL |
| 26 | 0.139 | 0.131 | 0.148 | 0.367 | EL |
| 32 | 0.238 | 0.202 | 0.274 | 0.367 | EL |
| 34 | 0.283 | 0.270 | 0.296 | 0.367 | EL |
| 36 | -0.200 | -0.200 | -0.200 | 0.367 | EL |
| 22 | -0.200 | -0.200 | -0.200 | 0.367 | MPM |
| 26 | 0.012 | -0.007 | 0.031 | 0.367 | MPM |
| 32 | 0.073 | -0.032 | 0.178 | 0.367 | MPM |
| 34 | 0.128 | 0.100 | 0.156 | 0.367 | MPM |
| 36 | -0.200 | -0.200 | -0.200 | 0.367 | MPM |
| 22 | 0.118 | 0.109 | 0.127 | 0.550 | AEL |
| 26 | 0.135 | 0.130 | 0.139 | 0.550 | AEL |
| 32 | 0.192 | 0.177 | 0.206 | 0.550 | AEL |
| 34 | 0.202 | 0.195 | 0.209 | 0.550 | AEL |
| 36 | -0.200 | -0.200 | -0.200 | 0.550 | AEL |
| 22 | 0.126 | 0.116 | 0.136 | 0.550 | EL |
| 26 | 0.178 | 0.170 | 0.185 | 0.550 | EL |
| 32 | 0.268 | 0.258 | 0.278 | 0.550 | EL |
| 34 | 0.238 | 0.210 | 0.266 | 0.550 | EL |
| 36 | -0.200 | -0.200 | -0.200 | 0.550 | EL |
| 22 | 0.010 | -0.064 | 0.084 | 0.550 | MPM |
| 26 | -0.002 | -0.024 | 0.020 | 0.550 | MPM |
| 32 | 0.080 | 0.033 | 0.128 | 0.550 | MPM |
| 34 | -0.042 | -0.075 | -0.008 | 0.550 | MPM |
| 36 | -0.200 | -0.200 | -0.200 | 0.550 | MPM |
| 22 | 0.119 | 0.112 | 0.125 | 0.733 | AEL |
| 26 | 0.156 | 0.143 | 0.168 | 0.733 | AEL |
| 32 | 0.181 | 0.163 | 0.199 | 0.733 | AEL |
| 34 | 0.198 | 0.192 | 0.203 | 0.733 | AEL |
| 36 | -0.200 | -0.200 | -0.200 | 0.733 | AEL |
| 22 | 0.134 | 0.120 | 0.148 | 0.733 | EL |
| 26 | 0.226 | 0.191 | 0.262 | 0.733 | EL |
| 32 | 0.247 | 0.225 | 0.270 | 0.733 | EL |
| 34 | 0.175 | 0.141 | 0.209 | 0.733 | EL |
| 36 | -0.200 | -0.200 | -0.200 | 0.733 | EL |
| 22 | -0.054 | -0.098 | -0.011 | 0.733 | MPM |
| 26 | 0.055 | 0.028 | 0.081 | 0.733 | MPM |
| 32 | 0.074 | -0.012 | 0.159 | 0.733 | MPM |
| 34 | -0.138 | -0.204 | -0.072 | 0.733 | MPM |
| 36 | -0.200 | -0.200 | -0.200 | 0.733 | MPM |

**Table S3:** *r*_m_ calculations (mean *±* standard deviation) calculated using the analytic *r*_m_ model (AE-L), the E-L equation, and the MPM for the larval competition dataset (Huxley et al., 2022) under the assumption that juvenile survival only decreased at the juvenile-to-adult transition.. Resource levels are in mg ml*^−^*^1^. Replicate-level *r*_m_ calculations are provided in an accompanying data file.

| temperature | estimate | lower | upper | resource level | model |
| --- | --- | --- | --- | --- | --- |
| 22 | 0.072 | 0.069 | 0.074 | 0.183 | AEL |
| 26 | 0.105 | 0.102 | 0.109 | 0.183 | AEL |
| 32 | 0.125 | 0.112 | 0.137 | 0.183 | AEL |
| 34 | 0.192 | 0.188 | 0.196 | 0.183 | AEL |
| 36 | -0.200 | -0.200 | -0.200 | 0.183 | AEL |
| 22 | 0.023 | 0.016 | 0.030 | 0.183 | EL |
| 26 | 0.011 | -0.023 | 0.045 | 0.183 | EL |
| 32 | 0.063 | 0.056 | 0.071 | 0.183 | EL |
| 34 | 0.132 | 0.084 | 0.179 | 0.183 | EL |
| 36 | -0.200 | -0.200 | -0.200 | 0.183 | EL |
| 22 | -0.027 | -0.036 | -0.018 | 0.183 | MPM |
| 26 | -0.042 | -0.110 | 0.027 | 0.183 | MPM |
| 32 | -0.028 | -0.095 | 0.038 | 0.183 | MPM |
| 34 | 0.112 | 0.087 | 0.137 | 0.183 | MPM |
| 36 | -0.200 | -0.200 | -0.200 | 0.183 | MPM |
| 22 | 0.099 | 0.096 | 0.102 | 0.367 | AEL |
| 26 | 0.130 | 0.126 | 0.134 | 0.367 | AEL |
| 32 | 0.184 | 0.161 | 0.207 | 0.367 | AEL |
| 34 | 0.243 | 0.222 | 0.264 | 0.367 | AEL |
| 36 | -0.200 | -0.200 | -0.200 | 0.367 | AEL |
| 22 | 0.067 | 0.065 | 0.069 | 0.367 | EL |
| 26 | 0.139 | 0.131 | 0.148 | 0.367 | EL |
| 32 | 0.238 | 0.202 | 0.274 | 0.367 | EL |
| 34 | 0.283 | 0.270 | 0.296 | 0.367 | EL |
| 36 | -0.200 | -0.200 | -0.200 | 0.367 | EL |
| 22 | 0.025 | 0.022 | 0.029 | 0.367 | MPM |
| 26 | 0.101 | 0.097 | 0.105 | 0.367 | MPM |
| 32 | 0.178 | 0.129 | 0.228 | 0.367 | MPM |
| 34 | 0.248 | 0.232 | 0.264 | 0.367 | MPM |
| 36 | -0.200 | -0.200 | -0.200 | 0.367 | MPM |
| 22 | 0.118 | 0.109 | 0.127 | 0.550 | AEL |
| 26 | 0.135 | 0.130 | 0.139 | 0.550 | AEL |
| 32 | 0.192 | 0.177 | 0.206 | 0.550 | AEL |
| 34 | 0.202 | 0.195 | 0.209 | 0.550 | AEL |
| 36 | -0.200 | -0.200 | -0.200 | 0.550 | AEL |
| 22 | 0.126 | 0.116 | 0.136 | 0.550 | EL |
| 26 | 0.178 | 0.170 | 0.185 | 0.550 | EL |
| 32 | 0.268 | 0.258 | 0.278 | 0.550 | EL |
| 34 | 0.238 | 0.210 | 0.266 | 0.550 | EL |
| 36 | -0.200 | -0.200 | -0.200 | 0.550 | EL |
| 22 | 0.093 | 0.071 | 0.114 | 0.550 | MPM |
| 26 | 0.115 | 0.112 | 0.117 | 0.550 | MPM |
| 32 | 0.200 | 0.180 | 0.221 | 0.550 | MPM |
| 34 | 0.152 | 0.119 | 0.185 | 0.550 | MPM |
| 36 | -0.200 | -0.200 | -0.200 | 0.550 | MPM |
| 22 | 0.119 | 0.112 | 0.125 | 0.733 | AEL |
| 26 | 0.156 | 0.143 | 0.168 | 0.733 | AEL |
| 32 | 0.181 | 0.163 | 0.199 | 0.733 | AEL |
| 34 | 0.198 | 0.192 | 0.203 | 0.733 | AEL |
| 36 | -0.200 | -0.200 | -0.200 | 0.733 | AEL |
| 22 | 0.134 | 0.120 | 0.148 | 0.733 | EL |
| 26 | 0.226 | 0.191 | 0.262 | 0.733 | EL |
| 32 | 0.247 | 0.225 | 0.270 | 0.733 | EL |
| 34 | 0.175 | 0.141 | 0.209 | 0.733 | EL |
| 36 | -0.200 | -0.200 | -0.200 | 0.733 | EL |
| 22 | 0.088 | 0.079 | 0.097 | 0.733 | MPM |
| 26 | 0.165 | 0.138 | 0.192 | 0.733 | MPM |
| 32 | 0.182 | 0.146 | 0.219 | 0.733 | MPM |
| 34 | 0.138 | 0.126 | 0.151 | 0.733 | MPM |
| 36 | -0.200 | -0.200 | -0.200 | 0.733 | MPM |

**Table S4:** Comparison of predicted *T*_opt_ and *r*_m_ at *T*_opt_ (*r*_opt_) with *±* bootstrapped 95% confidence intervals for the analytic *r*_m_ model, the E-L equation and the MPM. Letters “e” and “t” in the survival column indicate whether juvenile survival decreased at a fixed rate per time step or only at the juvenile-adult transition point, respectively. For the AE-L, juvenile survival can only decrease at the juvenile-to-adult transition (i.e., “t”).

| model | resource level | survival | TPC parameter | estimate | lower | upper |
| --- | --- | --- | --- | --- | --- | --- |
| AE-L | 0.183 | t | ropt | 0.17 | 0.15 | 0.19 |
| AE-L | 0.183 | t | Topt | 33.63 | 33.24 | 34.02 |
| E-L | 0.183 | e | ropt | 0.13 | 0.10 | 0.16 |
| E-L | 0.183 | e | Topt | 33.93 | 33.15 | 34.44 |
| E-L | 0.183 | t | ropt | 0.13 | 0.10 | 0.15 |
| E-L | 0.183 | t | Topt | 33.93 | 33.15 | 34.48 |
| MPM | 0.183 | t | ropt | 0.11 | 0.08 | 0.13 |
| MPM | 0.183 | t | Topt | 33.78 | 33.19 | 34.34 |
| AE-L | 0.367 | t | ropt | 0.23 | 0.21 | 0.25 |
| AE-L | 0.367 | t | Topt | 33.56 | 33.22 | 33.95 |
| E-L | 0.367 | e | ropt | 0.29 | 0.27 | 0.31 |
| E-L | 0.367 | e | Topt | 33.53 | 33.26 | 33.82 |
| E-L | 0.367 | t | ropt | 0.29 | 0.27 | 0.31 |
| E-L | 0.367 | t | Topt | 33.53 | 33.30 | 33.85 |
| MPM | 0.367 | e | ropt | 0.13 | 0.11 | 0.16 |
| MPM | 0.367 | e | Topt | 33.41 | 32.56 | 34.13 |
| MPM | 0.367 | t | ropt | 0.24 | 0.21 | 0.26 |
| MPM | 0.367 | t | Topt | 33.70 | 33.34 | 34.07 |
| AE-L | 0.550 | t | ropt | 0.21 | 0.20 | 0.22 |
| AE-L | 0.550 | t | Topt | 33.28 | 33.14 | 33.54 |
| E-L | 0.550 | e | ropt | 0.28 | 0.27 | 0.31 |
| E-L | 0.550 | e | Topt | 33.04 | 32.94 | 33.19 |
| E-L | 0.550 | t | ropt | 0.28 | 0.27 | 0.31 |
| E-L | 0.550 | t | Topt | 33.04 | 32.95 | 33.23 |
| MPM | 0.550 | e | ropt | 0.07 | 0.05 | 0.09 |
| MPM | 0.550 | e | Topt | 31.09 | 28.98 | 32.52 |
| MPM | 0.550 | t | ropt | 0.21 | 0.19 | 0.23 |
| MPM | 0.550 | t | Topt | 32.87 | 32.73 | 33.10 |
| AE-L | 0.733 | t | ropt | 0.20 | 0.19 | 0.21 |
| AE-L | 0.733 | t | Topt | 33.24 | 33.04 | 33.56 |
| E-L | 0.733 | e | ropt | 0.26 | 0.24 | 0.31 |
| E-L | 0.733 | e | Topt | 32.59 | 32.40 | 32.90 |
| E-L | 0.733 | t | ropt | 0.26 | 0.24 | 0.31 |
| E-L | 0.733 | t | Topt | 32.59 | 32.43 | 32.93 |
| MPM | 0.733 | e | ropt | 0.09 | 0.05 | 0.13 |
| MPM | 0.733 | e | Topt | 28.32 | 25.68 | 30.71 |
| MPM | 0.733 | t | ropt | 0.19 | 0.17 | 0.24 |
| MPM | 0.733 | t | Topt | 32.70 | 32.52 | 33.33 |

**Table S5:**
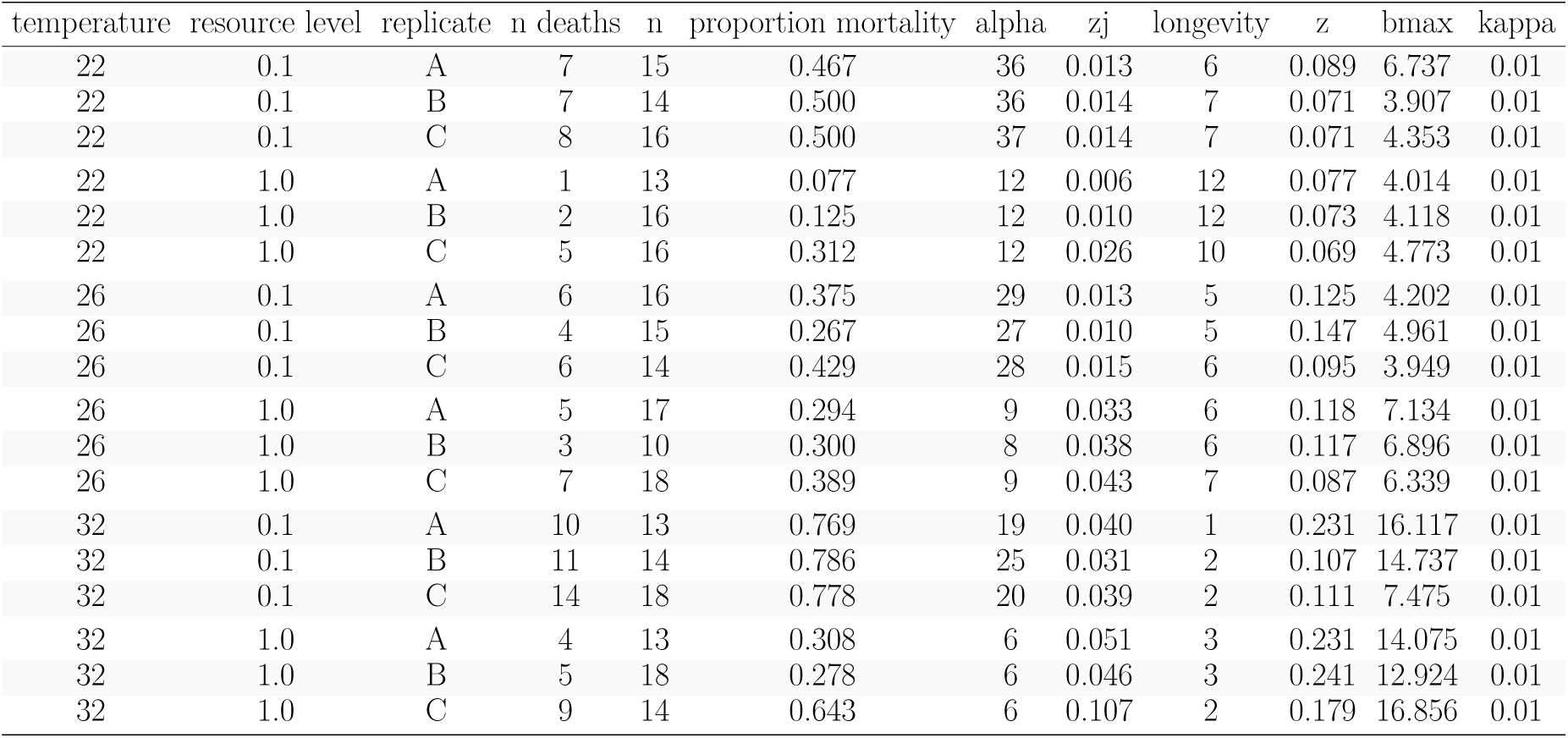
Replicate-level fitness trait values (means) used to calculate *r*_m_ using the analytic *r*_m_ model, the E-L equation, and the MPM for the resource limitation dataset (Huxley et al., 2021). *n* is the number of individuals in each replicate at the start of the experiment, *α* is egg-to-adult development time (days), *b*_max_ is peak reproductive rate (individuals (eggs) *×* individuals (females) *×* day^-1^), *κ* is the fecundity loss schedule (individual^-1^ day^-1^), and *z*_J_ and *z* are juvenile and adult mortality rates (individual^-1^ day^-1^), respectively.

**Table S6:** *r*_m_ calculations (mean *±* standard deviation) calculated using the analytic *r*_m_ model (AE-L), the E-L equation, and the MPM for the resource limitation dataset (Huxley et al., 2021). Resource levels are in mg larva*^−^*^1^ day*^−^*^1^. Letters “e” and “t” in the survival column indicate whether juvenile survival decreased at a fixed rate per time step or only at the juvenile-adult transition point, respectively. For the AE-L, juvenile survival can only decrease at the juvenile-to-adult transition (i.e., “t”). Replicate-level *r*_m_ calculations are provided in an accompanying data file.

| temperature | resource level | model | survival | estimate | lower | upper |
| --- | --- | --- | --- | --- | --- | --- |
| 22 | 0.1 | AEL | t | 0.074 | 0.067 | 0.080 |
| 26 | 0.1 | AEL | t | 0.088 | 0.082 | 0.095 |
| 32 | 0.1 | AEL | t | 0.129 | 0.112 | 0.146 |
| 22 | 0.1 | EL | e | 0.051 | 0.045 | 0.058 |
| 26 | 0.1 | EL | e | 0.062 | 0.053 | 0.070 |
| 32 | 0.1 | EL | e | -0.074 | -0.211 | 0.063 |
| 22 | 0.1 | EL | t | 0.051 | 0.045 | 0.058 |
| 26 | 0.1 | EL | t | 0.062 | 0.053 | 0.070 |
| 32 | 0.1 | EL | t | -0.074 | -0.211 | 0.063 |
| 22 | 0.1 | MPM | e | -0.239 | -0.262 | -0.217 |
| 26 | 0.1 | MPM | e | -0.101 | -0.143 | -0.058 |
| 32 | 0.1 | MPM | e | -0.460 | -0.487 | -0.432 |
| 22 | 0.1 | MPM | t | 0.037 | 0.028 | 0.046 |
| 26 | 0.1 | MPM | t | 0.083 | 0.040 | 0.127 |
| 32 | 0.1 | MPM | t | 0.047 | 0.029 | 0.066 |
| 22 | 1.0 | AEL | t | 0.157 | 0.154 | 0.160 |
| 26 | 1.0 | AEL | t | 0.217 | 0.198 | 0.235 |
| 32 | 1.0 | AEL | t | 0.359 | 0.343 | 0.376 |
| 22 | 1.0 | EL | e | 0.199 | 0.190 | 0.208 |
| 26 | 1.0 | EL | e | 0.252 | 0.233 | 0.271 |
| 32 | 1.0 | EL | e | 0.300 | 0.199 | 0.401 |
| 22 | 1.0 | EL | t | 0.199 | 0.190 | 0.208 |
| 26 | 1.0 | EL | t | 0.252 | 0.233 | 0.271 |
| 32 | 1.0 | EL | t | 0.300 | 0.199 | 0.401 |
| 22 | 1.0 | MPM | e | 0.091 | 0.017 | 0.165 |
| 26 | 1.0 | MPM | e | 0.078 | 0.028 | 0.128 |
| 32 | 1.0 | MPM | e | 0.176 | 0.024 | 0.329 |
| 22 | 1.0 | MPM | t | 0.163 | 0.145 | 0.180 |
| 26 | 1.0 | MPM | t | 0.211 | 0.187 | 0.235 |
| 32 | 1.0 | MPM | t | 0.337 | 0.283 | 0.390 |

